# The CcpNmr Analysis Simulated Metabolomics Database (CASMDB): An Open-Source Collection of Metabolite Annotation Data for 1D ^1^H NMR-Based Metabolomics

**DOI:** 10.1101/2024.05.05.592402

**Authors:** Morgan W. Hayward, Luca G. Mureddu, Gary Thompson, Marie Phelan, Edward J. Brooksbank, Geerten W. Vuister

## Abstract

Databases are invaluable for the identification of individual metabolites in untargeted metabolomics analyses, providing annotated pure metabolite references that allow for comparisons with experimentally collected mixture samples. Despite the value of an extensive reference database, publicly available databases for NMR-based metabolomics are often incomplete with respect to experimental conditions and derived NMR annotation parameters, such as peak positions. Hence, they are not designed for visualising the reference spectra alongside an experimental sample spectrum of interest, thus limiting the usefulness of the database. As a consequence, researchers have resorted to their own user- or application based database implementations.

In this paper we describe the collection, remediation and integration of annotation data from the publicly available HMDB, BRMB and GISMO NMR metabolomics databases to build the CcpNmr Analysis Simulated Metabolomics Database (CASMDB) that contains 1932 unique metabolite entries. This database, in concert with the AnalysisMetabolomics programme, also allows or accurate simulation of spectra at arbitrary field strengths. Together, these tools underpin the visualising of experimental and simulated metabolite references and their usage in 1D ^1^H NMR-based metabolomics studies.

## Background

Metabolomics is the scientific field of global systems biology concerned with small molecules, i.e. with molecular mass < 1.5KDa, that are substrates or products of metabolism, aka metabolites. The metabolites in tissue, fluids or biological samples in general constitute the metabolic profile. Changes in metabolite concentrations in such as a metabolic profile are examined typically downstream of the other ‘-omics’ i.e. genomics, transcriptomics and proteomics, and tend to be more influenced by external factors such as diet and disease (Wishart, 2019). As such, a metabolomic profile is a particularly good representative of the phenotype of the sample under examination. Metabolomics studies typically aim to identify biomarkers of metabolic processes or disease and have applications in a wide variety of research interest including pharmaceuticals (Alarcon-Barrera *et al*., 2022) personalised medicine (Di Minno *et al*., 2022), nutrition (LeVatte *et al*., 2021) microbiology (Zhao *et al*., 2020), agriculture (Razzaq *et al*., 2022), marine biology (Bayona, de Voogd and Choi, 2022) and toxicology (Dehghani *et al*., 2022).

The two premiere experimental techniques used in metabolomics analyses are Mass Spectrometry (MS) and Nuclear Magnetic Resonance (NMR) with each technique associated with its own advantages. MS has the greater sensitivity of the two techniques, making it favourable for identifying and quantifying metabolites with lower abundance in the sample or for precise quantification of a subset of known metabolites, also known as targeted metabolomics (Emwas, 2015). NMR-based metabolomics on the other hand, affords the inference of invaluable chemical knowledge albeit a lower intrinsic sensitivity. In addition, NMR-based metabolomics is highly automatable and reproducible with faster and simpler sample preparation compared to MS, making it a suitable technique for high-throughput procedures. In particularly, such procedures can be used for the observation and comparison of whole metabolite profiles without comprehensive (a priori) knowledge of the metabolites in the sample, also known as untargeted metabolomics (Emwas, 2015). For this paper, we focus on the database requirements, implementation and data that underpin NMR-based metabolomics, specifically one dimensional proton (1D-^1^H) NMR.

In NMR spectroscopy, a metabolite can be identified by a characteristic set of peaks in the NMR spectrum and each peak can be described by its chemical shift, i.e. position in the spectrum expressed in ppm, its signal intensity and its width at half height expressed in Hz. Metabolite NMR spectra can also be further described by assigning peaks to coupling patterns or ‘multiplets’ where the peaks in the multiplet represent the collective signal of the protons in a specific chemical environment. For example, the signal from a methyl group in an aliphatic compound may appear as three peaks with an intensity ratio of 1:2:1 and would be referred to as a triplet. In the same fashion that a metabolite signature is the collective signal of all its peaks, the spectra obtained in NMR-based metabolomics are the sum of the signatures of all metabolites in the sample and can often comprise hundreds to thousands of peaks. In order to discern biologically relevant information from the experimental metabolomics spectra of a biological sample, the experimentally observed peaks, must be assigned to their corresponding metabolite(s), typically by referencing to the previously measured spectra of metabolite standards. Therefore, high-quality databases of metabolite spectra, whether from publicly available sources or otherwise, are invaluable to NMR-based metabolomics researchers (Heyward & Vuister, 2023).

The two most well-known publicly accessible databases for NMR-based metabolomics are the Human Metabolome Data Base (HMDB) (Wishart, Guo *et al*., 2022) and the Biological Magnetic Resonance Data Bank (BMRB) (Hoch *et al*., 2022) (cf. Table 1). The HMDB contains entries for > 240,000 metabolites and contains extensive biological metadata including metabolite chemical details, typical concentrations in biological samples and ontological classification. However, HMDB is primarily focused on MS spectra rather than NMR, with many entries lacking experimental NMR spectra. The BMRB is an NMR-specific database containing > 2,700 experimentally supported entries making it particularly reliable for NMR-based researchers, but unfortunately lacks the extensive metabolite metadata of the HMDB.

**Table 1:**
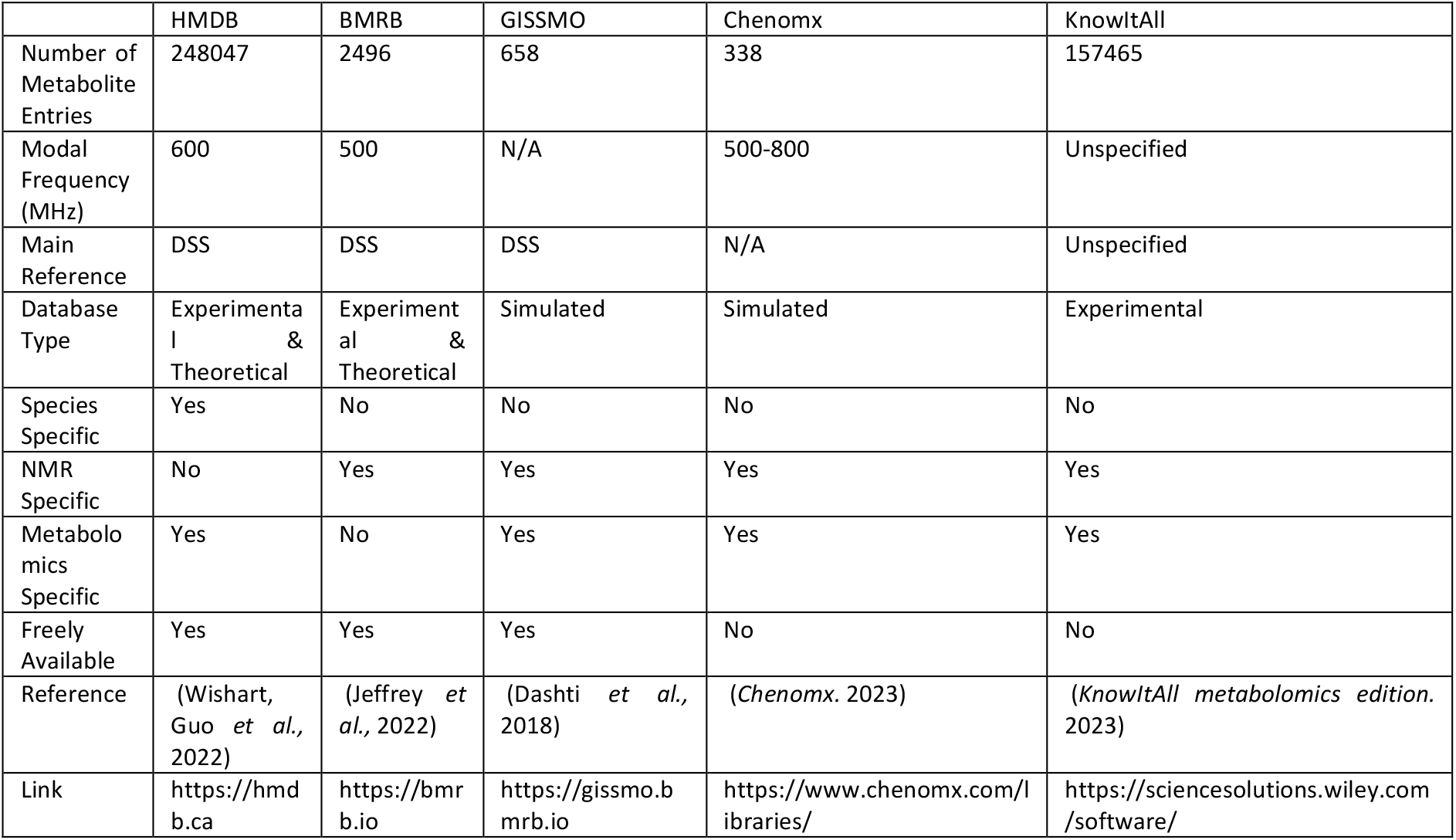
Overview of commonly used freely available and commercial databases for NMR-based metabolomics.

Commercial metabolomics databases (cf. Table 1) could also be an option for NMR-based metabolomics research. They typically come as an integrated part of an NMR analysis software package. The Chenomx NMR Suite (*Chenomx*. 2023) contains a database of annotated spectra for 336 compounds at a range of spectrometer frequencies and is particularly thorough on the effect of changing pH on the metabolite spectra. Alternatively, the KnowItAll Metabolomics Edition (*KnowItAll metabolomics edition*. 2023) comes with and extensive spectral database of ∼160,000 compounds, many of which are pharmaceuticals or drug screening fragments. In general, these databases tend to be more consistent in terms of sample conditions and annotation compared to the publicly available counterparts, but require costly subscriptions. Moreover, their closed-source nature often results in reduced usage flexibility, making them even more difficult to justify for academic research projects.

Given that metabolite NMR spectra can vary depending on the sample and spectral conditions, e.g. pH, solvent, temperature and spectrometer field strength, the ideal database would need a reference for each metabolite for every value of every condition for all possible spectrometers. Creating such an encompassing database would be a herculean task and well beyond the scope of most research groups. Alternatively, with good understanding of metabolite physio-chemical properties and NMR theory, a thoroughly annotated spectrum with its respective metabolite chemical data can be simulated at any condition or field strength. The Guided Ideographic Spin System Model Optimization library (GISSMO) (Dashti *et al*., 2018) is an example of one such effort; it contains the information of metabolite spectra abstracted as spin system matrices (SSMs) that encode the chemical shift and coupling constants of the protons in a given metabolite. Peak-lists generated from these SSMs are remarkably accurate and are completely field independent, meaning one entry can account for all potential spectrometer frequencies. This is notably only feasible for small to medium metabolites as the simulation algorithm has an exponential time complexity.

The CcpNmr AnalysisMetabolomics program is part of a software suite for biomolecular NMR that also includes AnalysisAssign (Skinner *et al*., 2016) for general NMR data analysis and AnalysisScreen (Mureddu *et al*., 2020) for NMR-based small molecule screening. As a part of our effort to develop AnalysisMetabolomics, we gathered metabolite spectra annotation data from the HMDB, BMRB and GISSMO libraries to build a freely available database of NMR-based metabolomics spectra annotation data, denoted as CcpNmr AnalysisMetabolomics Database (CASMDB) for simulating reference spectra for spectral deconvolution purposes.

## Methods

Annotation data, i.e. spectral peak lists and metabolite meta data were downloaded from the HMDB (https://hmdb.ca), BMRB (https://bmrb.io/metabolomics) and GISSMO (https://gissmo.bmrb.io) repositories for all available experimentally entries in various formats (*vide infra*). Entries that are not supported by experimental data or are not annotated completely were labelled appropriately for future awareness. Theoretical datasets, i.e. peak-lists based off of chemical structure alone, were not included in this database. Table 1 gives an overview of relevant statistics.

### HMDB Files

HMDB annotation data for NMR-based metabolomics is non-inclusively available across a range of different formats and so all formats were downloaded from the website (https://hmdb.ca/downloads) for parsing to ensure complete data coverage. All metabolite biological and chemical data was available as a single large Extensible Markup Language (XML) file, whereas sample and spectral data, including peak-lists, were available in XML format (n=2924), text format (TXT) (n=1002) and/or NMR Markup Language format (nmrML) (n=773). Where appropriate, files were refined to allow automatic parsing; see supplementary Table 1 for a full description of file refinement and file exclusions.

Metabolite data was extracted automatically in a priority system where nmrML files were parsed first due to their inclusion of supporting spectral arrays and multiplet assignment, followed by XML files and then TXT to supplement sample and spectrum data whenever nmrML data only proved insufficient. When matching data across multiple files, the HMDB accession numbers were used to match metabolites to their annotation data and HMDB experiment numbers were used to match sample, spectrum and peak data. Peak width data from the HMDB was mostly unavailable and so an arbitrary width of 1Hz was used as a placeholder in order to accommodate simulation requirements.

### BMRB Files

BMRB experimental data were retrieved in bulk from the BMRB website (https://bmrb.io/metabolomics/) in Self-Defining Text Archive and Retrieval (STAR) file (n=2496) format. These original data were parsed using the official BMRB API python package (Wedell, 2022) to retrieve metabolite, sample and spectral data, including peak data, however not multiplet assignments as this information is unavailable in the BMRB repository. Metabolite meta data was limited to name, description, chemical formula, average molecular weight, Simplified Molecular-Input Line-Entry System (SMILES) and International Chemical Identifier number (InChI). Of the 2496 files, 1414 files were excluded due to insufficient peak annotation data, see Supplementary Table 1 for details. Similarly to the HMDB, peak width data was unavailable, so a placeholder of 1 Hz was used.

### GISSMO Files

GISSMO spin system matrices were downloaded from the GISSMO website (https://gissmo.bmrb.io) in XML format (n=659). Spin system matrices were extracted automatically and stored in tabular format; proton chemical shifts were stored in the multiplets table and coupling constants in the coupling table which contains the coupling values and the proton/multiplet identifiers for coupling assignment. As GISSMO entries are an extension of BMRB entries, metabolite, sample and spectral data are inferred from BMRB data with the same accession numbers. No peak data was available from the GISSMO files due to the nature of the data.

### Data Verification and Remediation

To ensure that the NMR spectrum simulations are representative they were compared against their respective experimental spectra where data availability allowed such a comparison. Simulated spectra were generated to exactly match the experimental spectra with respect to sweep width and number of data points. Simulation accuracy was assessed using an unscaled dot-product, also known as cosine similarity, which scores between 1 for a perfect match and -1 for a perfect inverted result. Overall similarity scores were calculated from the mean of the similarity scores of spectrum sub-sections containing simulation peak data, i.e. the regions of interest, to prevent solvent and reference peaks interfering with the similarity scores. A total similarity score of 0.9 was used as a threshold of sufficient accuracy of the simulated spectrum. All simulations with similarity scores below 0.9 were automatically adjusted for overall peak widths and alignments. Any simulations that could not be automatically modified to reach a similarity score of >= 0.9 were visually assessed and remediated manually.

Simulations were remediated by updating peak- and multiplet data in peak-list based simulations and by adding spin-spin couplings in spin-system based simulations. Appropriate peak and multiplet data were gathered by peak picking and multiplet assignment of the associated experimental spectra in the CcpNmr AnalysisMetabolomics programme, which is derived from the AnalysisAssign software (Skinner et al., 2016). All remediations were made with reference to the chemical structure of the sample metabolite to ensure appropriate multiplet assignment. All experimental spectra were imported into AnalysisMetabolomics using their native binary formats, but converted and saved in Hierarchal Data Format (HDF5) files (to be published elsewhere) for user reference, including any processes such as phase correction and chemical shift reference alignments.

### Database Layout

All data processed from the respective databases was saved into a tabular format for compact storage and efficient access when remediating. The finalised database was converted to a directory of NMR Exchange Format (NEF) files, with each NEF file documenting the data for an unique metabolite as to allow ease of expansion and exclusion of metabolite data for users. The database was made with a structure that reflects the biological relationships of the data as well as the data structures of the CcpNmr AnalysisMetabolomics programme to allow easy access and deposition to and from the software (Fig. 1). Due to the abundance of metadata from the HMDB, the core metabolites table and the metabolite metadata tables were structured similarly to the HMDB metabocards. However, the ontology data from these metabocards is stored in single table as opposed to the hierarchical format in the HMDB. Metabolites from the BMRB and GISSMO repositories were matched to HMDB entries via International Chemical Identifier (InChI) and stored under the same entry in the metabolites table to avoid duplicates. Unique CASMDB IDs were created for each table to ensure proper distinct data relationships. Database origin accession numbers were stored in the most appropriate tables, e.g. HMDB accessions are unique and unambiguous at the metabolite level whereas BMRB accessions are ambiguous at the metabolite level, therefore HMDB and BMRB accessions were stored in metabolite and sample tables, respectively.

**Figure 1.**
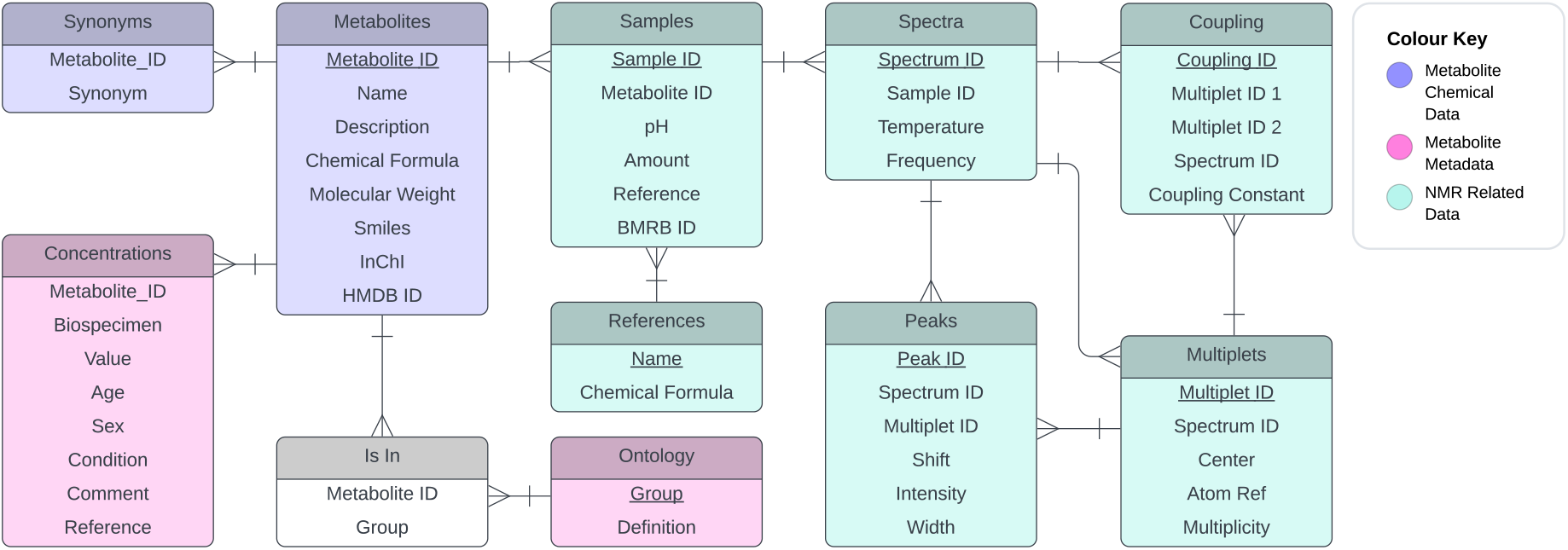
Entity Relationship (ER) diagram depicting the structure of the CcpNmr Metabolomics Database and the relationships between the various tables. Colours indicate classification of the data contained in the table: metabolite data (purple), metabolite meta data (pink), NMR data (teal). Connectors indicate the relationships between the tables, all of which are many- to-one, e.g. a metabolite entry is linked to many samples, but each sample is linked to one metabolite. Ontology is the only exception where multiple metabolites can be in multiple biological samples, hence the inclusion of the ‘Is In’ table to create a many- to-many relationship.

To access and deposit data into the database, a user interface module was developed to allow user-controlled extraction of data from the database for simulating reference spectra. Database entries are transformed into CcpNmr data objects of the equivalent type, e.g. sample data to CcpNmr sample objects and metabolite data to CcpNmr substance objects.

### Spectrum Simulation

Spectrum simulations were created using modified functions using the publicly available nmrsim (Sametez, 2021) python package for simulating the NMR spectra of spin-1/2 nuclei. Simulated spectra are created from the metabolite data by transforming peak chemical shifts, heights and widths into Lorentzian distributions which are summed to create the spectrum intensity arrays. For entries without peak lists, i.e. GISSMO entries, peak lists are created from multiplet chemical shift and coupling constants via the nmrsim quantum mechanics (qm) functions. A second user interface was developed in the CcpNmr Analysis V3 Metabolomics software to allow users to select metabolites of interest at available sample and spectrum conditions and, where multiplet assignment is present, to shift multiplets to account for signal drift. Subsequently, a fully annotated 1D-1H spectrum in the CcpNmr metabolomics software can be automatically transformed into a user-defined database entry. Details of the user interfaces will be reported elsewhere.

## Results

Overall, 1932 unique metabolite entries, as determined by their InChI codes, were included in CASMDB, with 3329 samples, 3333 spectra, 24021 multiplets and 71485 peaks (Table 2). The HMDB was the highest contributor of metabolite entries of the three databases queried; it is also the only contributor for ontology and concentration data. Although the relationship from sample to spectrum is considered as one-to-many, i.e. there can be multiple spectra recorded for a single sample, this was only the case in 4 samples with the vast majority of entries only having one 1D 1H spectrum for each sample. No multiplets were extracted from the BMRB entries as peak to atom assignment was not possible from star files. No peak data was imported from GISSMO entries due to the nature of the data.

**Table 2:** Summary of the content of the CcpNmr Metabolomics database. Note that the total for metabolites and synonyms is not the sum of the corresponding numbers of the individual databases, as the databases are not mutually exclusive.

|  | HMDB | BMRB | GISSMO | Merged Total |
| --- | --- | --- | --- | --- |
| Metabolites | 1372 | 972 | 581 | 1932 |
| Synonyms | 40031 | 12865 | 0 | 52566 |
| Ontology | 53731 | 0 | 0 | 53731 |
| Concentrations | 24250 | 0 | 0 | 24250 |
| Samples | 1471 | 1200 | 658 | 3329 |
| Spectra | 1471 | 1204 | 658 | 3333 |
| Multiplets | 19201 | 0 | 4820 | 24021 |
| Peaks | 38931 | 32554 | 0 | 71485 |

Metabolite distribution across the databases was greater than expected with only 458 of the 1932 metabolites shared between the HMDB and the BMRB repositories, when considering the GISSMO entries as extensions of the BMRB (Fig. 2). Even though GISSMO entries were developed from BMRB data, data for forty-six metabolites were extracted from the GISSMO repository only, but somewhat surprisingly not from the BMRB. This is most likely due to the many exclusions required for the initial parsing of the BMRB data (cf. Supplementary Table 1) due to missing peak data. The data retrieved from the HMDB was the most exclusive with 914 metabolites not found in either of the other databases.

**Figure 2.**
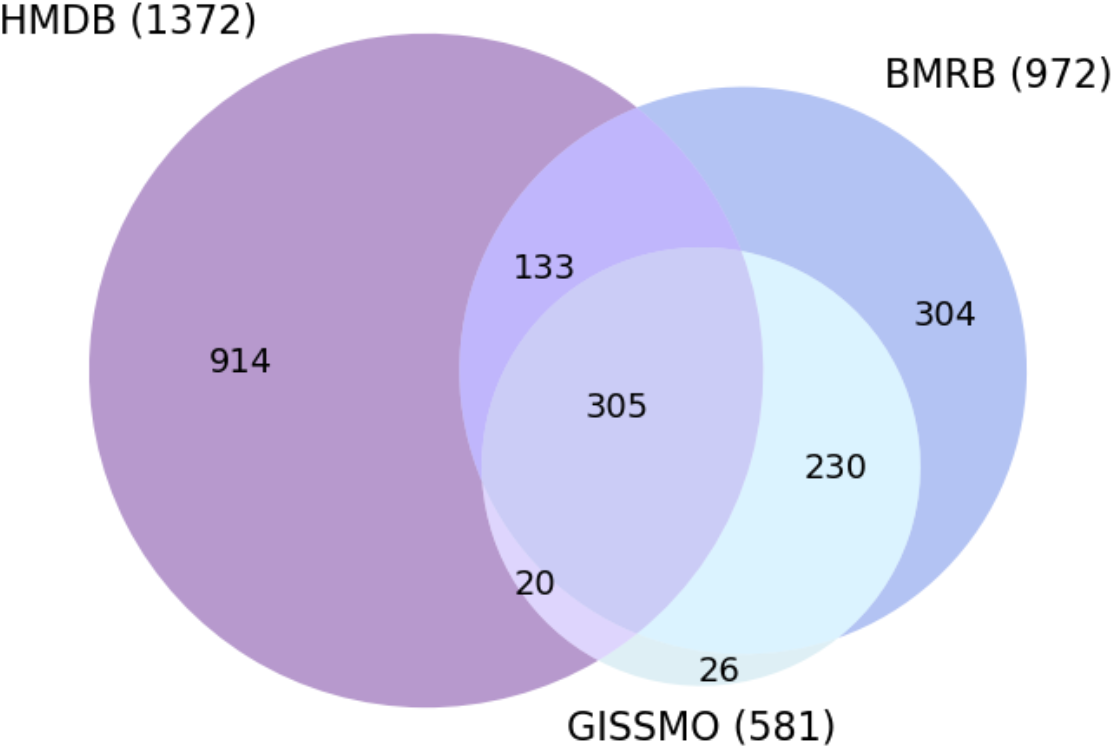
Venn diagram showing the CASMDB metabolite distribution as derived from the source databases. Numbers indicate unique metabolite counts. Metabolites were identified as identical across databases on the basis of their InChI codes.

There was a range of sample and spectral conditions for the data from each source, with the HMDB data typically showing the largest variation (Fig. 3). Most samples were at a pH between 6.5 and 7.5, with HMDB spectra typically recorded at pH 7.0 (n=855) and BMRB at pH 7.4 (n=691). The full range of pH values was between 1 and 12 for HMDB samples and between 7 and 12 for BMRB samples. There were many samples with missing pH values for both HMDB and BMRB data. Sample references were most commonly DSS or TMS, with BMRB samples exclusively one of these references. A minority of HMDB samples used CHCl_3_ or TSP as the reference compound. Six samples had missing values for reference compounds. The spectrometer frequency ranges for the HMDB spectra were between 90 and 700 MHz, whereas the BMRB spectra were exclusively recorded at 400, 500 and 600 MHz. Fifty-one HMDB entries had missing data for the spectrometer frequency, whereas only one BMRB spectrum (for entry bmse000060) was missing this data. Temperature data are not displayed in Fig. 3 as all spectral data were recorded at 298K.

**Figure 3:**
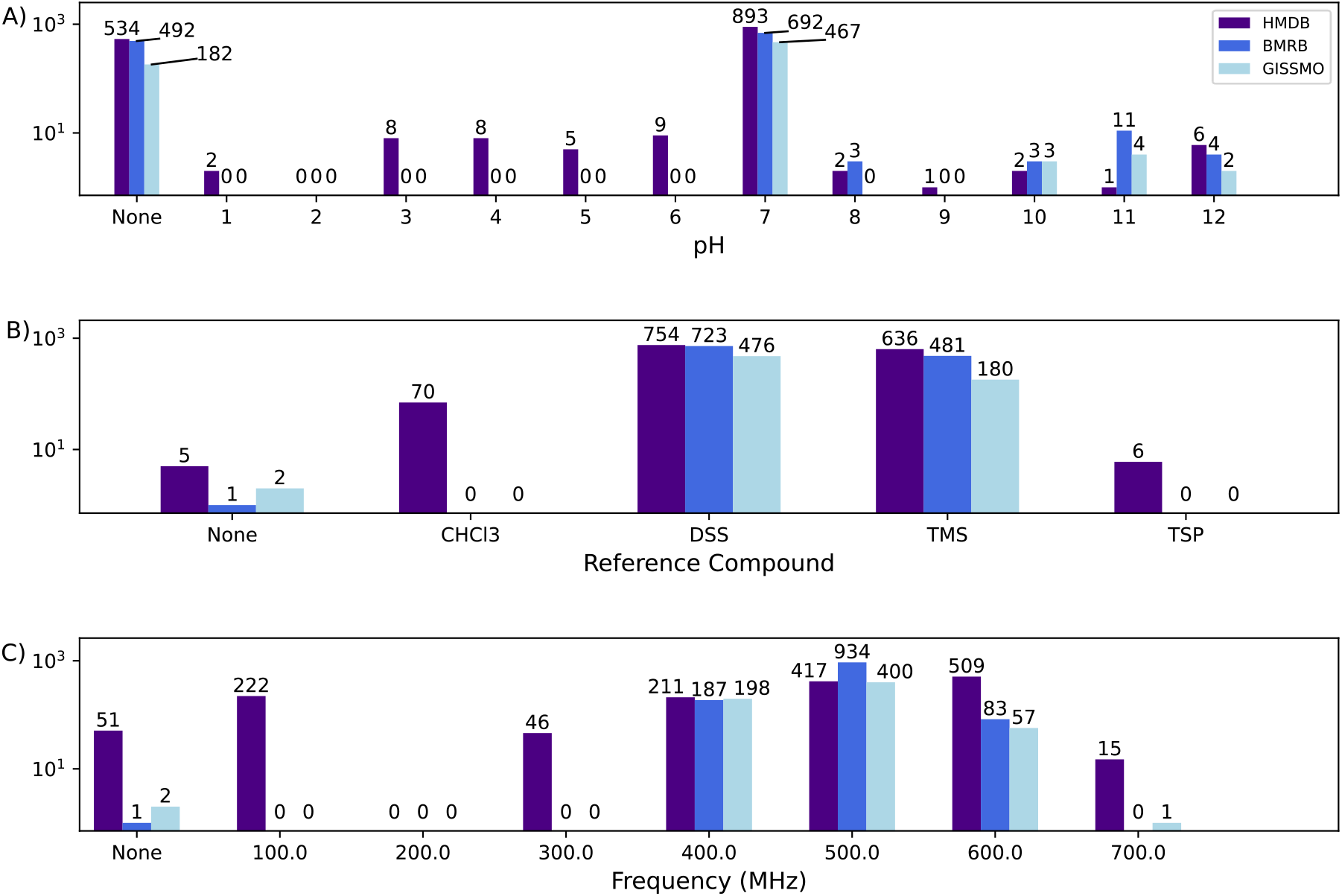
Experimental conditions of the CASMDB entries colour coded according to source. Count numbers are shown above the bars and all Y axes are in logarithmic scale. A) Bar chart depicting the distribution of pH values in the samples, rounded to nearest integer values. B) Bar chart depicting the spectral referencing compound used. C) Bar chart depicting the distribution of spectrometer frequencies in the metabolite spectra, rounded to the nearest 100 MHz. None indicates the entries with an un-specified value.

Simulated spectra could be recreated from peak-list data, to act as a handle for identification of potential issues and relative quantification purposes. Fig. 4 shows an example of simulated ^1^H NMR spectra of L-Valine based on peak and multiplet data derived from the three databases, compared to an experimental ^1^H NMR spectrum. In this example, all simulations are visually representative of the real spectrum and can be easily matched despite the relatively high signal to noise ratio of the experimental spectrum and the subtle differences in peaks in the simulated spectra. HMDB and BMRB simulations are made from annotated peak-lists whereas GISSMO simulations generate peak-lists based on coupling interactions. This can be seen in the examples where the GISSMO simulation appreciates the smaller peaks at the edges of the multiplet that would be lost in the noise.

**Figure 4.**
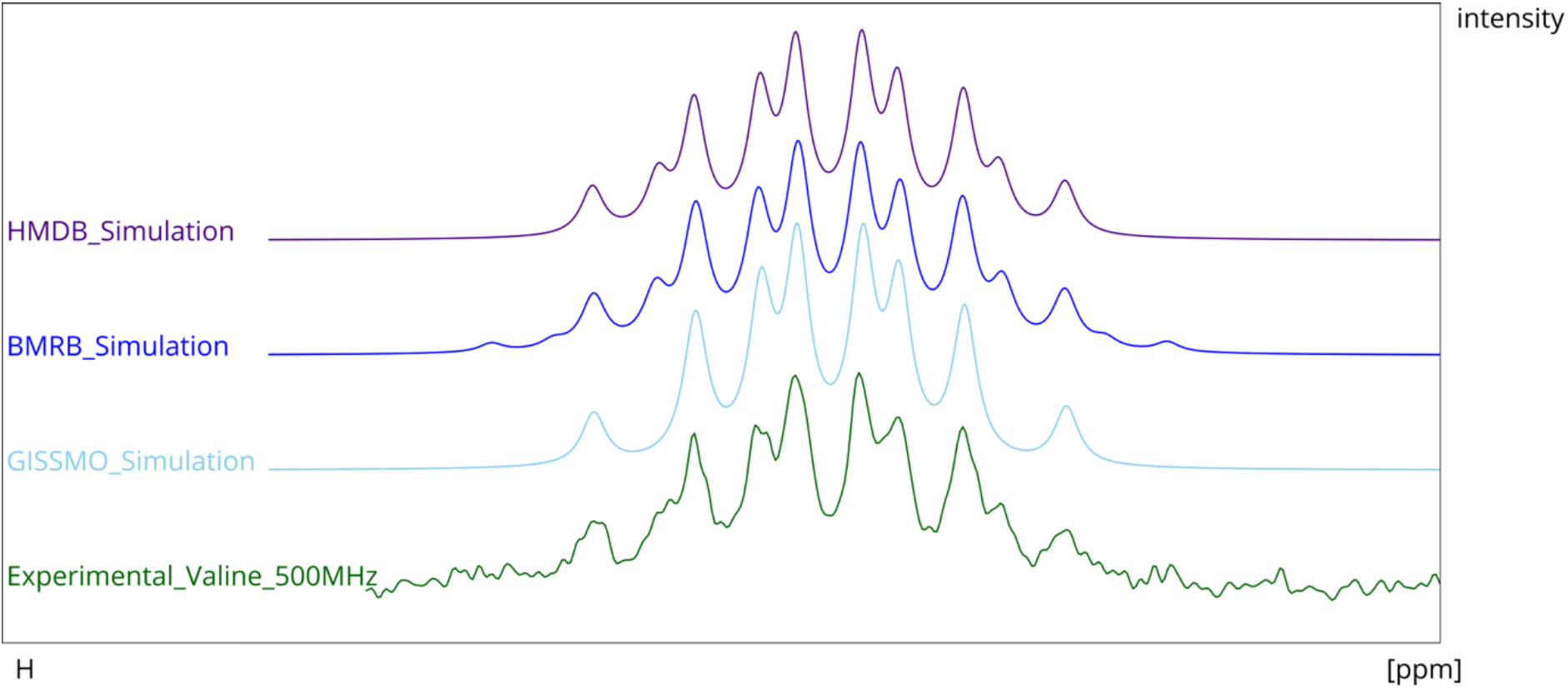
Comparison of an experimental ^1^ H NMR spectrum to simulated spectra based on peak- and multiplet data contained in the various databases. The plots show the multiplet s ignal at 2 .28 ppm of the L-Valine H^β^ - proton. The experimental spectrum shown was retrieved from the BMRB, accession number: BMSE 000052. The HMDB s imulated spectrum was shifted by ∼ 0. 03 ppm to align to the other spectra.

Nevertheless, all the simulations are still representative to distinguish a complex splitting pattern. It is noteworthy that simulated spectra will lack common features of experimental spectra, such as noise, artefacts, reference peaks and water/solvent signals. Such simulations also do not include any errors in the line shape of the spectra, such as resulting from poor phasing or non-Lorentzian line shapes due to poor shimming.

A total of 1469 simulated spectra had to undergo remediations to reach a suitable level of accuracy, of which 510 manual remediations were made to peaks/couplings (Fig. 5A). Most manual remediation was made to the HMDB data, most often as result of low precision in peak positions. Many HMDB spectra were recorded at a precision of +-0.005ppm, which is typically not enough to recreate multiplet patterns with sufficient precision, as many peaks become indistinguishable from each other. The primary remediation requirement for BMRB data was to increase the overall peak width, often to 3Hz. Only five entries from the GISSMO database required adding of additional couplings, with most simulations being highly representative of the actual experimental data. A small subset of poorly matching simulations could not be remediated due to either low quality or incorrect spectra in the experimental file. Any simulation that could not be verified, including simulations without any associated experimental spectra, are still included in the database but marked as unverified. Supplementary Fig. 1 shows typical examples of remediation and exclusion. Figure 6 shows the overall improvement of simulation accuracy after remediation.

**Figure 5:**
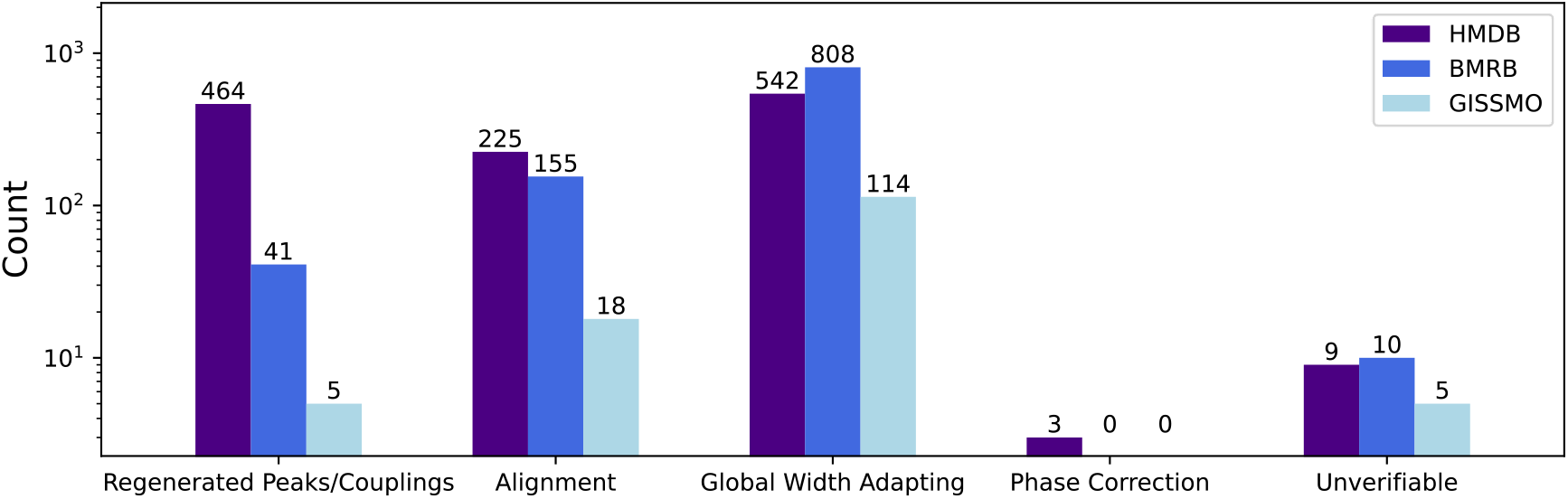
Bar chart of the remediation categories, separated by database of origin. Count number refers to the number of entries that were remediated. Note the logarithmic Y-scale.

**Figure 6.**
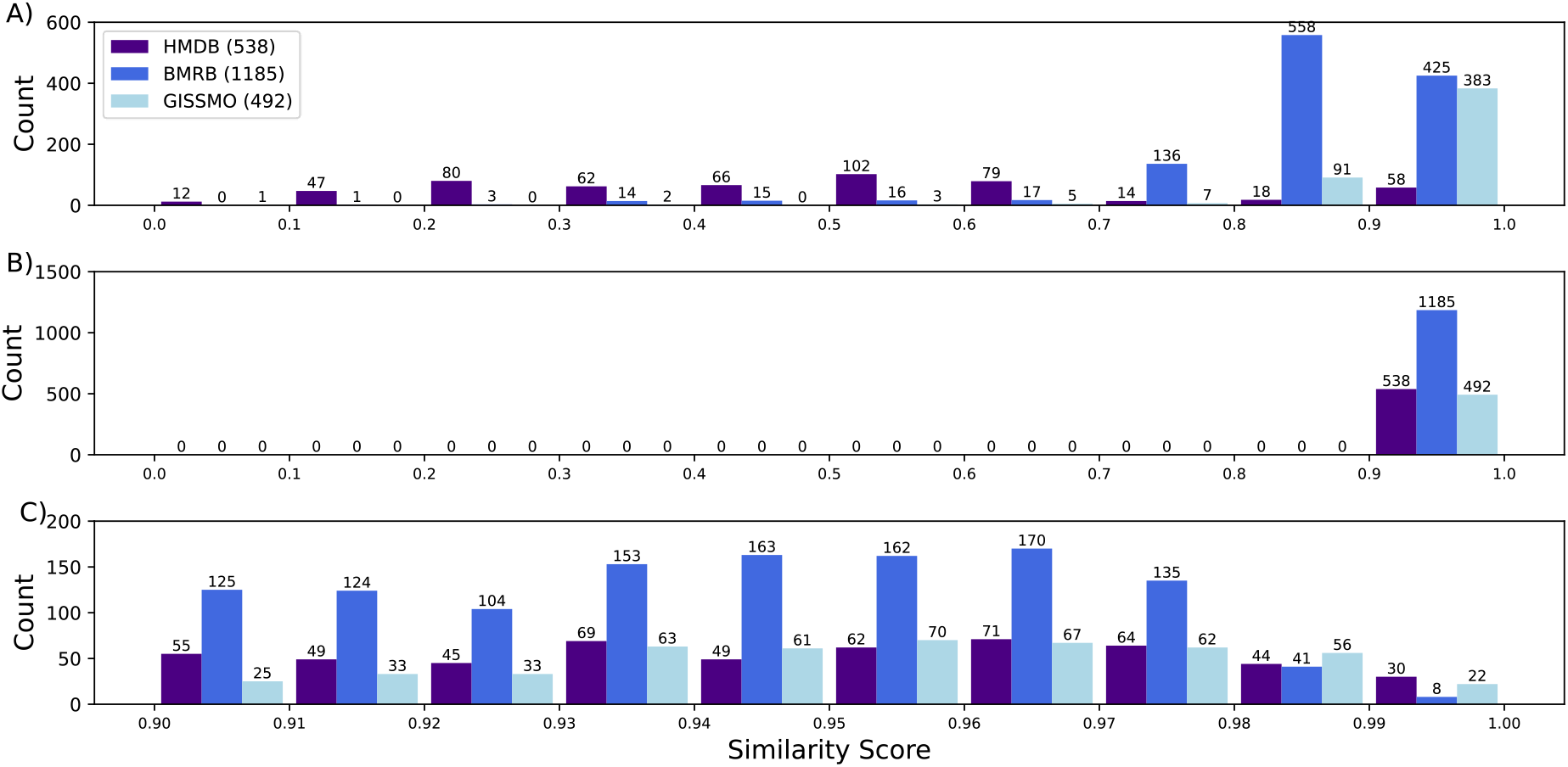
Histogram of the cosine similarity accuracy scores before (A) and after (B, C) remediations. C) Expanded scores from B).

## Discussion

### Database Structure

The structure of the database was built with minimalism and simplicity in mind, as it was not intended to contain detailed chemical- or spectral information, but rather was intended as source for generating metabolite spectra for identification and quantification purposes. As such, this database was not intended and not likely to replace publicly available databases like the HMDB and BMRB, nor function as a storage mechanism of the spectral simulation results. Instead, it was to aid the flexible, real-time visualisation of any data found in these databases.

The core of the database comprises the data required to recreate spectra, but each CASMDB entry also contains the necessary IDs and accession numbers to locate the original source. This simplicity also ensures that the database is relatively compact, allowing easy downloading and thus rapid access to metabolite overlays. The use of peak-lists and spin-system matrices over spectrum intensity and position arrays also contribute to the compactness of the database, as a spectrum can be recreated from tens to hundreds of peak values rather than storing tens of thousands of values in the form of (binary) intensity arrays. A simpler database format is also advantageous for user interaction, as it allows quick and easy deposition of custom metabolite references.

The database also contains tables for crucial metabolite metadata, which is not strictly needed for accurate reference overlays, but was included for user convenience. Firstly, it allows quick reference to biological context of the molecule without having to trace back the entry to its respective database web-page. This also allows the database to operate without an internet connection. Secondly, it allows for additional annotations, such as ontologies, by the user that might currently be absent from the data. We envision such information to ultimately feedback into CASMDB to enhance its functionality for all users.

### Database Content

The total of 1932 distinct metabolites can be considered large from a practical perspective, as most NMR-based metabolomics studies will typically identify/quantify 50 to 200 metabolites, which is an order of magnitude less than the total in CASMDB. However, 1932 metabolites are surprisingly few considering the total number of metabolites available at the source databases, with the HMDB alone containing entries for > 240,000 distinct entries. This highlights an alarmingly large issue of missing annotated data in NMR-based metabolomics databases due to many contributing factors. One such factor may simply be the lack of data deposition; the HMDB is substantially more supportive toward MS-based metabolomics, with many entries containing dozens of MS spectra but only one, or often no NMR spectra. Although it may be reasonable to expect a greater number of depositions for the technique with greater sensitivity, recent improvements in NMR hardware and the improving access to high-field NMR spectrometry means resolution in NMR-based metabolomics is ever improving and database references will need to be expanding accordingly.

Another factor for missing annotation data is that public databases were likely never designed for simulation purposes but rather for searching and matching to peak-lists. This is most evident with the BMRB database which creates its peak-lists by automated peak-picking of experimental spectra, thus allowing users to search for hits via peak-lists. Furthermore, over half of the BMRB entries had to be excluded due to insufficient peak data (Supplementary Table 1), mostly due to lack of peak intensities which are not necessary for searching by peak-list, but vital for creating simulations.

Another potential issue with the scope of the database content concerns the sample and spectral conditions. Ideally, we would have every metabolite recorded at several relevant conditions, but this is rarely the case. Spectrum differences caused by minor variations in pH and temperature can often be accounted for by allowing some flexibility in the chemical shifts of multiplets, and even spectrometer frequency can be extrapolated by rescaling peak positions relative to the multiplet they are assigned to. These adaptations are only possible if peak to atom assignments are present, which is the case in the HMDB and GISSMO entries but not in the BMRB entries. The BMRB entries instead account for condition differences, namely spectrometer frequency differences, by having more spectra for the same metabolite, as shown by the higher metabolite to spectrum ratio in the BMRB entries.

### Spectrum Simulations

Visual inspection of simulated spectra show that those created from peak-lists only are highly suitable as references, particularly for manual identification purposes and particularly for examining weak-coupling J-splitting patterns. The value of peak-list based simulations is reduced in more dense regions of the spectra where the peaks overlap considerably and where signal to noise is low. Under these circumstances, the contributions of the signals of individual metabolites start overlapping, rendering their exact deconvolution problematic. This effect is further compounded by low signal-to-noise ratios. Under these circumstances, proper spectrum similarity validation using other technologies is necessary; however this will be reported on in another study.

Spin-system based simulations do not suffer from this problem as they construct the simulations from first-principles, essentially starting with a single signal for each proton and distributing that signal throughout the splitting pattern resulting from the J-coupling interactions. This makes GISSMO simulations particularly accurate with minimal data input. Nevertheless, there are limitations for spin-system simulations as well. Firstly, they require accurate measurements for multiplet chemical shift and J-coupling values, something that can be particularly difficult in more complex spectra and take longer to record and analyse compared to automated peak-picking. Secondly, for medium to large metabolites, spin-system simulations take significantly longer to generate computationally than spectra produced from peak-lists only as the algorithm to produce peak-lists from spin-system matrices has an exponential time complexity. This means spin system simulations are best suited to very small metabolites i.e. with 10 or fewer coupled protons, or else they cannot be observed and manipulated in real time.

### Future Work

No database is complete upon its creation and CASMDB is no exception. CASMDB will need updating, expanding and maintaining. Expansion efforts include gathering additional metabolite spectra from other publicly available databases. Good candidates include the Natural Product Magnetic Resonance Database (NPMRD) (Wishart, Sayeeda *et al*., 2022), an NMR-specific database of small molecules developed by the Wishart Lab in a similar style to the HMDB although not specific to human metabolites. A second database is the Birmingham Metabolite Library (BML) (Ludwig *et al*., 2012), a relatively small but particularly thorough database of 1D ^1^H and 2D J-resolved NMR spectra of metabolites under a range of pH values and acquisition times. Extending CASMDB to include more spectra allows for a wider coverage and therefore increased usefulness for the metabolomics community, particularly when addressing metabolic questions that are non-human centred. Furthermore, in the case of duplicate entries, spectra at multiple conditions can be collected to yield information regarding relevant experimental factors or increase confidence in the spectra from identical duplicates.

An alternative to simply collecting and annotating more spectra is presented here where we develop and expand the use of spin systems and spectral simulations in NMR-based metabolomics. Simulations are exceptionally flexible by nature and so theoretically spectra only need to be recorded once to extract the parameters to cover all sample and spectral conditions. With modern NMR spectrum viewing software, proper annotation of spectra is easier than ever, including the creation and refinement of simulations from spin system matrices. We plan to develop an CcpNmr AnalysisMetabolomics module for real time creation of spin system matrices from measured experimental metabolite standards, thus allowing user creation of spin-system based simulations and entry into CASMDB.

### Final Remarks

The CASMDB functions as an free and easy-access reference database of simulating representative metabolite spectra for NMR-based metabolomics research. It has been built with the explicit purpose of spectral simulations and consequently has a concise structure dedicated to storing only the necessary parameters for simulated spectra. We expect and hope the database to grow in due course, as well as receive additional tools for spectral manipulation and custom reference creation.

## Acknowledgements

We thank all authors and contributors to the HMDB, BMRB and GIZMO databases. We also thank the members of the CCPN development team, Vicky Higman and Eliza Ploskon, for valuable discussion and feedback. MH is supported by a Leicester Institute of Structural and Chemical Biology PhD studentship. The research was supported by UKRI MRC grant MR/V000950/1 (to GWV).

## Abbreviations

CASMDB: CcpNmr Analysis Simulated Metabolomics Database
NMR: Nuclear Magnetic Resonance
MS: Mass Spectrometry
HMDB: Human Metabolome Database
BMRB: Biological Magnetic Resonance Data Bank
GISSMO: Guided Ideographic Spin System Model Optimization
SSMs: Spin System Matrices
1D ^1^H: one Dimensional proton
SQL: Structured Query Language
SMILES: Simplified Molecular-Input Line-Entry System
InChI: International Chemical Identifier number

## Notes

### Competing Interest Statement

The authors have declared no competing interest.

