## Supplementary materials for "The CcpNmr Analysis Simulated Metabolomics Database (CASMDB): An Open-Source Collection of Metabolite Annotation Data for 1D ^1^H NMR-Based Metabolomics"

### Supplementary Data

Morgan W. Hayward[1], Luca G. Mureddu[1], Gary Thompson[2], Marie Phelan[3] , Edward J. Brooksbank[1] and Geerten W. Vuister[1,*]

[1] Department of Molecular and Cell Biology, Leicester Institute of Structural and Chemical Biology, University of Leicester, Henry Wellcome Building, Lancaster Road, Leicester LE1 7HN, United Kingdom

[2] School of Biosciences, Division of Natural Sciences, University of Kent, Canterbury, CT2 7NZ, United Kingdom

[3] Institute of Systems, Molecular and Integrative Biology, NMR Centre for Structural Biology, University of Liverpool, Crown Street, Liverpool, L69 7ZB, United Kingdom

* Corresponding author

| Data Source | File Count | File Type | Refinement or Exclusion | Details |
|---|---|---|---|---|
| HMDB | 208 | nmrML | Refinement | Adding ‘/’ to the closing statement on line 5 and replacing ‘</chemicalShiftStandard>’ with ‘/>’ at the closing statement on line 11 |
| | 4 | nmrML | Refinement | Removing the ‘>’ character from the opening statement |
| | 3 | nmrML | Refinement | Removing the ‘<’ from the value string |
| | 1 | nmrML | Refinement | Replaced ‘-‘ with ‘Nan’ to allow conversion into float |
| | 5 | nmrML | Exclusion | No values for peaks or multiplets |
| | 12 | TXT | Refinement | Added ‘\t’ to allow division of data |
| | 11 | TXT | Refinement | Added ‘-‘ to empty space to recognise Null values |
| | 16 | TXT | Refinement | Swapped assignment and multiplet table titles for correct recognition |
| | 1 | TXT | Refinement | Removed a rogue character (‘v’) from file |
| | 2 | TXT | Refinement | Added a missing column title |
| | 1 | TXT | Refinement | Added a new line character after a table title |
| | 1 | TXT | Refinement | Removed an empty column with no title |
| BMRB | 1 | STR | Refinement | Edited filename for real spectrum. |
| | 600 | STR | Exclusion | No spectral peak list or assigned chemical shift |
| | 46 | STR | Exclusion | Unsuitable peak chemical shift data |
| | 768 | STR | Exclusion | Unsuitable peak intensity data |
| GISSMO | 6 | XML | Refinement | Spin system label changed to ‘merged’. |

Supplementary Table 1: A record of refinement and exclusion details of metabolite reference files from the BMRB and HMDB.

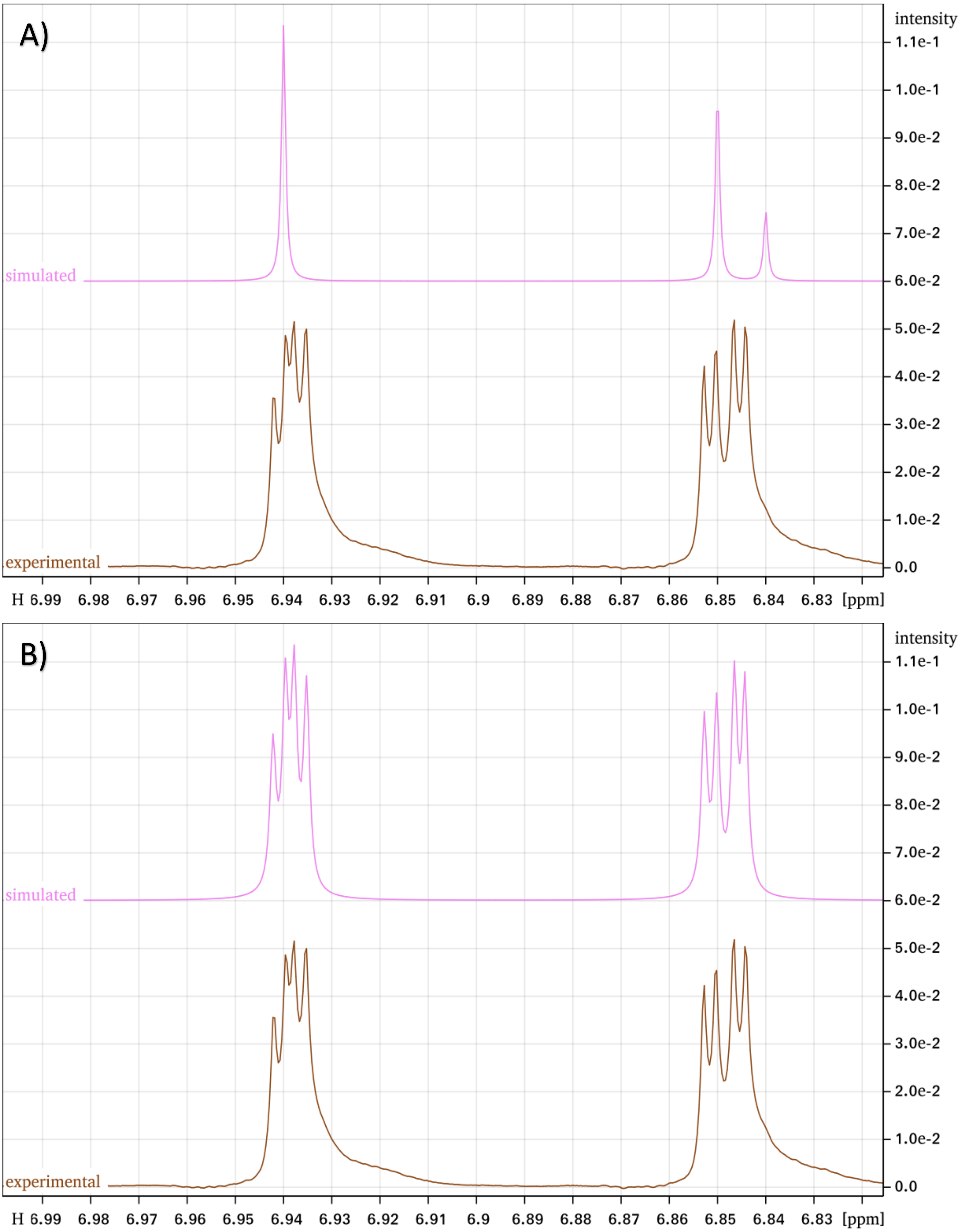


Supplementary Figure 1.1: Example of low peak precision in HMDB spectra before (A) and after (B) manual remediation of peak list peak chemical shift, height and width. Peaks with chemical shift recorded to a precision of +-0.005ppm could not be used to recreate accurate spectral line-shapes.

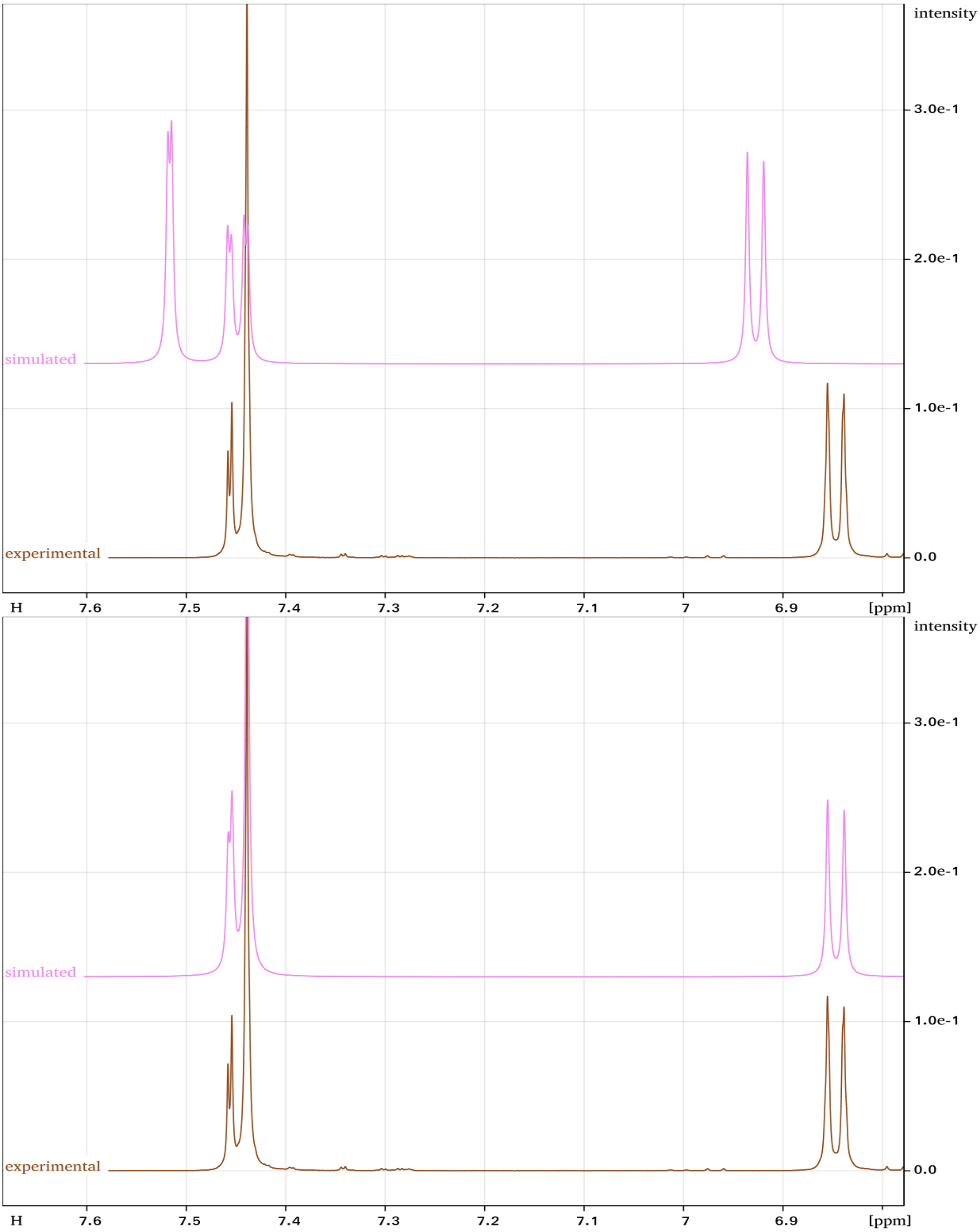


Supplementary Figure 1.2: Example of mild discrepancies between peak lists and the experimental spectra before (A) and after (B) manual remediation of peak list peak chemical shift, height and width.

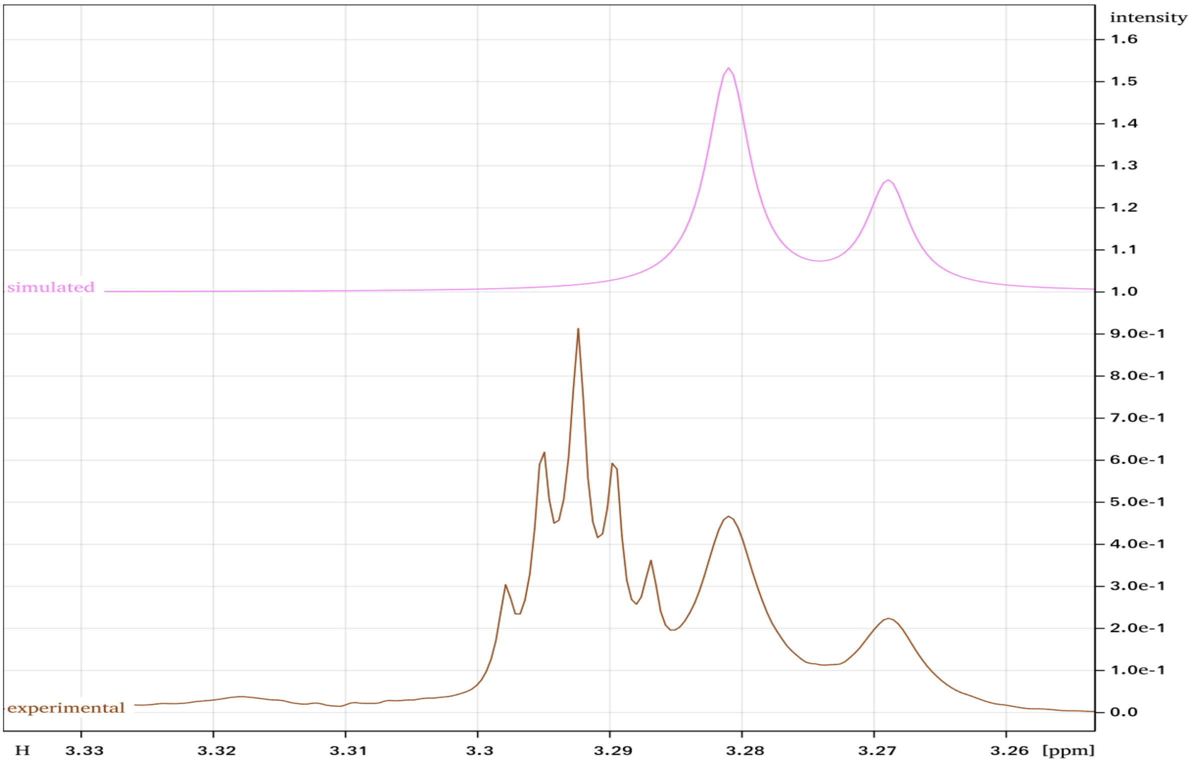


Supplementary Figure 1.3: Example of interference in the metabolite signal by the solvent signal. In this example, the Deuterated methanol quintuplet is close enough to the metabolite signal to effect the similarity scoring.

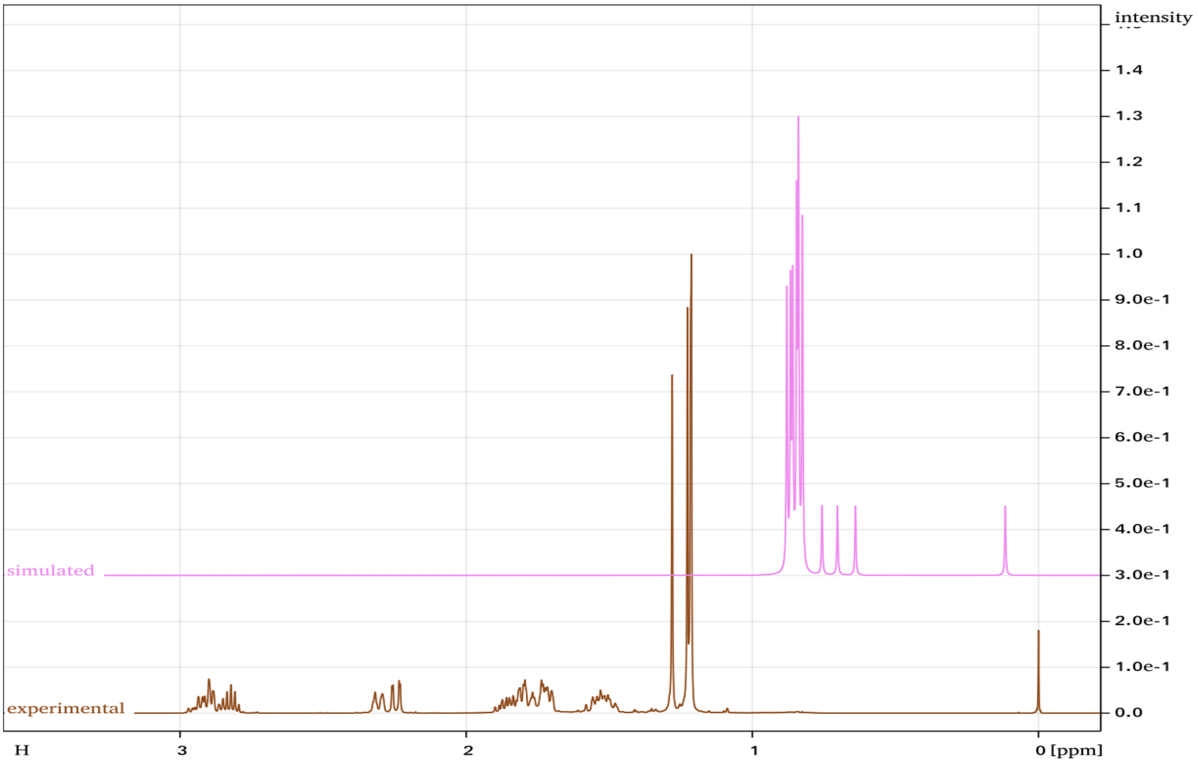


Supplementary Figure 1.4: Example of an exclusion due to extreme differences between experimental and simulated spectra.
